# msmu: a computational framework for provenance-aware mass spectrometry-based proteomics analysis

**DOI:** 10.64898/2026.01.07.698308

**Authors:** Hyung-Wook Choi, Byeongchan Lee, Un-Beom Kang, Sunghyun Huh

## Abstract

Heterogeneous data formats and analysis pipelines in computational proteomics hinder reproducibility, interoperability, and reuse of processed data. We present msmu, an open-source Python package that implements a modular, provenance-aware pipeline for post-search processing. Within a single MuData object, msmu links representations across peptide–spectrum match, peptide, and protein levels to machine-actionable provenance that records operations, parameter choices, software versions, and the execution environment. The object’s provenance log directly enables workflow replay with content-hash verification, without separately maintained scripts or provenance records. We demonstrate cross-platform reproducibility between macOS and Debian; replay reproduces the original protein inference map exactly (Jaccard indices = 1), with protein q-values and intensities agreeing to machine precision (root-mean-square deviation ≤ 2.0×10^−17^; maximum absolute difference ≤ 1.8×10^−15^). Furthermore, the same data structure supports seamless integration of proteomics with other MuData-compatible modalities. By coupling proteomics data with verifiable provenance in a single object, msmu advances FAIR-aligned reuse and multimodal interoperability.

## Main

Advances in bioinformatics tools have driven the wide adoption of mass spectrometry (MS)-based bottom-up proteomics in biological research, yet its computational landscape remains fragmented with less standardized data structures and processing pipelines compared to other omics disciplines^1,2^. As a result, reproducibility and reuse have become persistent challenges in proteomics. Reanalysis of public proteomics datasets often fails or yields divergent results^3–5^, as reproducing an analysis without recorded provenance amounts to a cumbersome reconstruction from scattered files, code, and prose^4,5^. Despite increasing awareness and FAIR (Findable, Accessible, Interoperable, and Reusable)-oriented guidelines on how reproducibility and reuse depend critically on factors such as transparent metadata, open data formats, and provenance that records how results were produced, these requirements remain inconsistently implemented^1,6,7^. Much of this gap arises downstream of database search, where peptide– spectrum matches (PSMs) are transformed and aggregated into peptide- and protein-level representations. These steps are often implemented through disjointed workflows, proprietary “black box” software, or *ad hoc* scripts. Although existing data formats can represent hierarchical proteomics data, there are few data models that couple data representations with machine-actionable provenance and can support workflow replay and verification. Collectively, these practices limit the reproducibility, interoperability, and reuse of proteomics data, and can impede downstream innovation that would benefit from greater computational transparency. Existing open-source tools^8–14^ address individual elements of this problem but do not integrate post-search processing with provenance embedded in the data object that enables workflow replay and facilitates data reuse.

Here we present msmu, an open-source Python package that provides a framework for modular, provenance-aware analysis of proteomics data. By leveraging MuData^15^, a widely used data format for multimodal omics, msmu organizes PSM-, peptide-, and protein-level representations as separate AnnData^16^ objects within a single MuData object. Building on this data structure, msmu provides a flexible end-to-end post-search pipeline—from search engine output parsing through preprocessing to statistical analysis—with processing operations that automatically capture provenance and co-locate it with the data they transform. Importantly, unlike tools that adopt AnnData or MuData primarily as storage containers (Supplementary Table 1), msmu treats provenance as a first-class component of the data object that is machine-actionable through content-hash verification and workflow replay, with exportable workflow scripts and environment specifications. For each operation, msmu records the function call, parameter choices, software versions, execution environment, and parent relationships defining data lineage, together with content hashes of the corresponding input and output data states. These records are serialized together with the data in a single .h5mu file based on HDF5^17^. The resulting provenance log allows msmu to replay workflows and verify regenerated data states against their recorded content hashes, without separately maintained workflow scripts or an external provenance system.

msmu comprises four core modules spanning data ingestion, preprocessing, statistical analysis, and visualization (Fig. 1a). The Data Ingestion module converts heterogeneous search engine outputs— currently supporting PSM or precursor data from Sage^18^, DIA-NN^19^, MaxQuant^20^, FragPipe^21^, and DelPi^22^—into a standardized MuData representation, and sample metadata in the Sample and Data Relationship Format (SDRF) are ingested alongside the quantitative data. The Data Preprocessing module provides feature- and sample-level filtering, hierarchical summarization including feature selection and intensity aggregation, protein inference, normalization, and batch effect correction. The Data Analysis module supports sample correlation, dimensionality reduction, and differential expression (DE) analysis. The Visualization module provides interactive plots for quality control metrics and downstream analytical results.

**Fig. 1.**
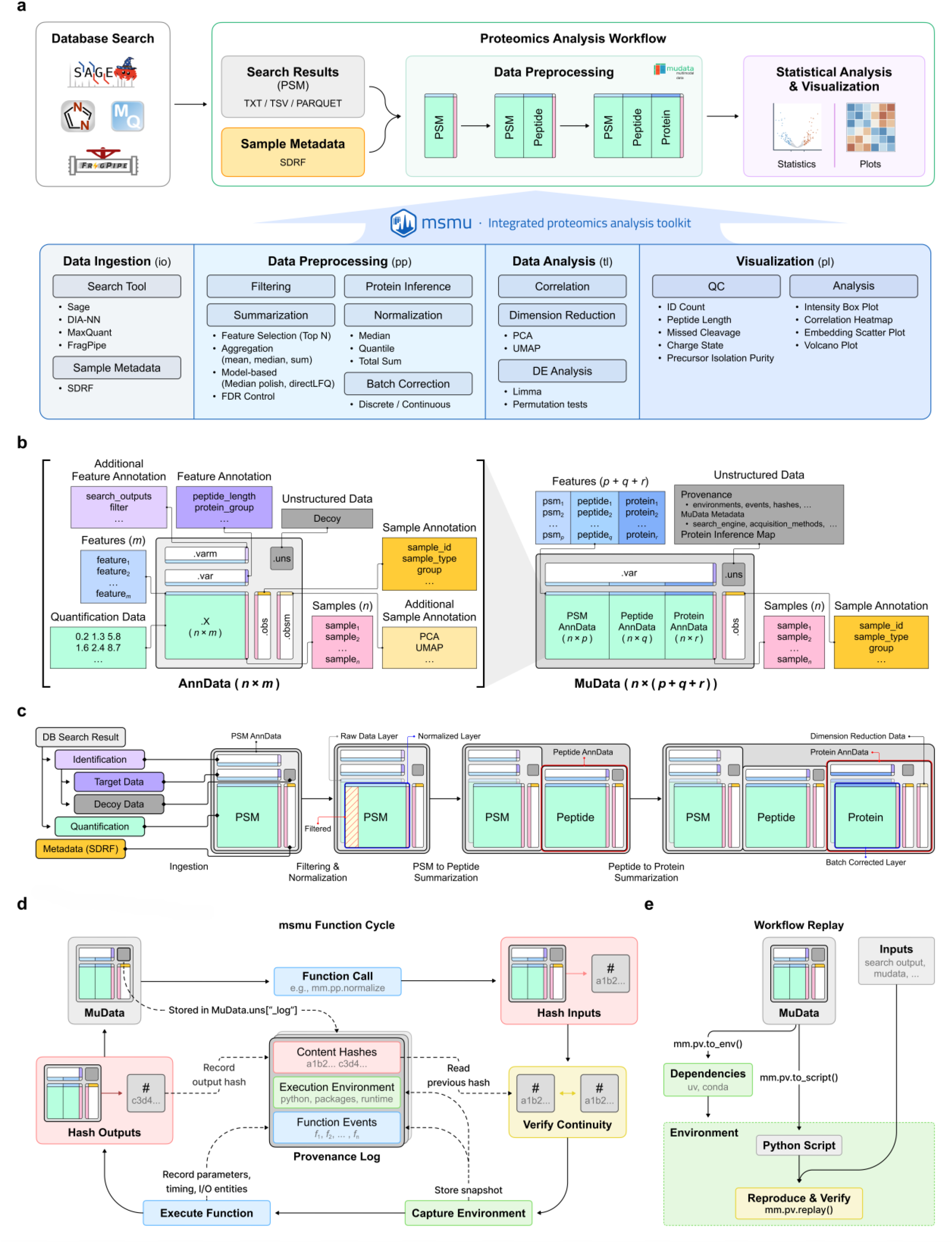
Architecture, data organization, provenance recording, and workflow replay of msmu. **a**, Overview of the msmu post-search analysis pipeline and its four modules for data ingestion (Data Ingestion), preprocessing (Data Preprocessing), statistical analysis (Data Analysis), and visualization (Visualization). PSM, peptide–spectrum match; FDR, false discovery rate; PCA, principal component analysis; UMAP, Uniform Manifold Approximation and Projection; DE, differential expression; QC, quality control; ID, identification; SDRF, Sample and Data Relationship Format. **b**, MuData organization used by msmu. The PSM-, peptide-, and protein-level data are represented as AnnData objects within a parent MuData object, which stores shared sample annotations, the provenance log, cross-level metadata, and the protein inference map. **c**, Example transformation of PSM-level search results into peptide- and protein-level representations. Filtering and normalization create distinct layers at the PSM level, while downstream analyses at the protein level, such as PCA, store derived representations in corresponding AnnData slots. DB, database. **d**, Function cycle. When a data-transforming function is called, its provenance decorator hashes the input data state, verifies continuity with the previously recorded output hash, captures the environment, executes the requested function, records the output hash in the MuData provenance log. **e**, Workflow replay. The provenance log can be exported as a Python script and environment specification, or used to replay the workflow from source to reproduce the original data.

Fig. 1b illustrates the hierarchical MuData organization used by msmu. Each data level is represented as an AnnData object containing quantitative matrices (.X), feature annotations (.var/.varm), sample annotations (.obs/.obsm), and auxiliary unstructured data (.uns). A parent MuData object wraps these AnnData objects and stores, in its global .uns, cross-level annotations, the provenance log (execution environments, recorded events, and content hashes), search engine metadata, and the protein inference map (peptide–protein graph). In a typical workflow (Fig. 1c), PSM-level search results and sample metadata are ingested into a PSM AnnData object, with the filtered and normalized matrix represented in separate data layers. Hierarchical summarization subsequently generates peptide- and protein-level AnnData representations within the same MuData object. Analysis functions store derived annotations and representations in corresponding AnnData slots. For example, protein-level principal component analysis (PCA) stores sample embeddings in .obsm and feature loadings in .varm, whereas batch effect correction adds a distinct layer containing the corrected protein matrix in the protein AnnData.

msmu stores provenance within the MuData object itself (Fig. 1d). When a data-transforming function is called, msmu hashes the input data states and verifies continuity against the previous output hash in the provenance log. msmu then captures the execution environment, executes the function while recording its parameters, timing, and input/output entities as a function event, and hashes the outputs, recording them as content hashes. Because each event references its parent events, the provenance log forms a graph of data lineage. These records are stored in MuData.uns[“_log”] and therefore travel with the .h5mu file. Functions that do not modify the MuData object (e.g., plotting functions) are not included in the provenance log. msmu implements workflow replay through functions that read the provenance log of a MuData object (Fig. 1e). mm.pv.to_env() exports the recorded Python and package versions as an environment specification for uv or conda, and mm.pv.to_script() exports an editable Python script listing every recorded call with its parameters and recorded hashes. mm.pv.replay() instead replays the workflow directly from the provenance log. Replay and the exported script both start from the source input files, rerun the recorded steps, verify each regenerated data state against its recorded content hash, and yield a new MuData object with its own provenance log. For customized reuse rather than reproduction, a script exported with verify=False omits these checks, so parameters, input files, or analysis steps can be edited, and the modified run is recorded anew (Methods). The provenance log thus supports both exact reproduction and customized reuse of prior data without separately maintained scripts or provenance records.

To evaluate msmu in a complete analysis workflow, we reanalyzed a previously published multi-omics dataset^23^ in which the proteome and transcriptome of murine epithelial (C10) and endothelial (SVEC) cells were profiled from the same single cells. The proteomics data were researched with Sage and quantified with FlashLFQ^24^, and the resulting PSM identification and quantification outputs were processed with msmu into a MuData object (Fig. 2a; Methods). Plots for protein identification per cell and cell type, correlation, and DE results can be readily generated using built-in functions in the Visualization module (Fig. 2b–e). To test whether the recorded provenance could reproduce processing operations and data states across computational environments, we exported the recorded environment from the .h5mu object originally generated on macOS and replayed the workflow on Debian. Because replay ran with hash verification enabled, its completion confirmed that every regenerated intermediate data state matched its recorded content hash under the 12-significant-digit normalization policy (Methods). Replay reproduced the protein inference map exactly, with a Jaccard index of 1 for 3,910 peptide nodes, 933 protein nodes, and 3,973 peptide–protein edges (Fig. 2f), so all 933 proteins rested on identical peptide evidence. Protein q-values were identical (root-mean-square deviation [RMSD] = 0; Fig. 2g), and log_2_ protein intensities differed only at floating-point scale (RMSD = 2.0×10^−17^; maximum absolute difference [max |Δ|] = 1.8×10^−15^; Fig. 2h). DE analysis repeated on the replay-derived protein matrix yielded q-values agreeing to within machine precision (RMSD = 2.4×10^−16^; max |Δ| = 5.3×10^−15^), and all significance calls were preserved at q < 0.10, 0.05, and 0.01 (274, 248, and 187 proteins, respectively; Fig. 2i). Together, these results show that provenance-driven replay in msmu preserves the numerical and inferential outputs of a recorded workflow.

**Fig. 2.**
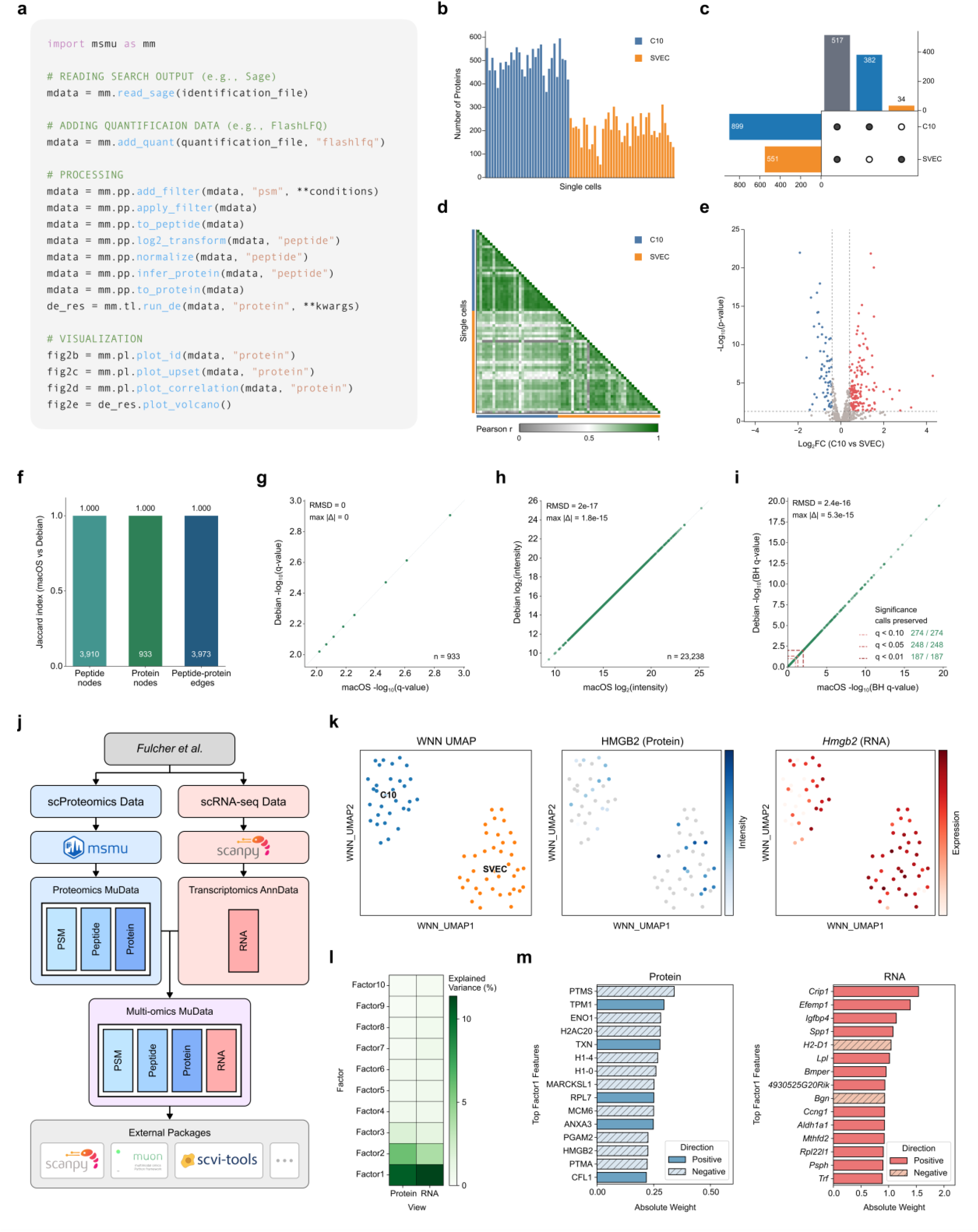
End-to-end proteomics analysis, cross-platform replay, and multi-omics integration with msmu. **a**, Example pseudocode for an msmu-based proteomics analysis workflow of a previously published dataset^23^. **b**, Number of proteins identified per single cell. **c**, UpSet plot of protein identified exclusively in or shared between C10 and SVEC cells. **d**, Pearson correlation heatmap of protein intensities between cells. **e**, Volcano plot of differential protein abundance between C10 and SVEC cells. **f**, Jaccard indices comparing peptide nodes, protein nodes, and peptide–protein edges between the original macOS analysis and the Debian replay. The numbers of nodes and edges are indicated inside the respective bars. **g**, Comparison of −log_10_ protein-level q-values between the original and replayed analyses. The number of proteins is indicated. RMSD, root-mean-square deviation; max |Δ|, maximum absolute difference. **h**, Comparison of log_2_ protein intensities between the original and replayed analyses. The number of non-missing protein–cell values is indicated. **i**, Comparison of BH-adjusted q-values (-log_10_) from differential expression (DE) analysis performed on the original and replay-derived protein matrices. The numbers of preserved significance calls (i.e., differentially expressed proteins) at q < 0.10, 0.05, and 0.01 are indicated. BH, Benjamini–Hochberg. **j**, Multimodal data integration workflow. The single-cell proteomics (scProteomics) and matched single-cell RNA-sequencing (scRNA-seq) data were processed using msmu and scanpy, respectively. Proteomics data were represented as a MuData object and scRNA-seq data were added as an additional AnnData modality, forming a single multimodal MuData object. **k**, WNN UMAP computed from the integrated data (left), and HMGB2 protein intensity (log_2_, middle) and *Hmgb2* RNA expression (natural log, right) visualized on the same UMAP embedding. WNN, Weighted Nearest Neighbor; UMAP, Uniform Manifold Approximation and Projection. **l**, Percentage of variance explained by each factor per modality in MOFA. MOFA, Multi-Omics Factor Analysis. **m**, Absolute MOFA factor weights for the top 15 protein (left) and RNA (right) features of Factor 1. Solid and hatched bars indicate positive and negative weights, respectively.

We next evaluated direct interoperability of the msmu-processed proteomics object with established multimodal analysis workflows. The matched single-cell RNA-sequencing (scRNA-seq) data were processed with scanpy^25^ into an AnnData object (Fig. 2j; Methods). The proteomic and transcriptomic representations were combined into a multimodal MuData object, enabling direct use of scverse tools such as scanpy and muon^15^. Weighted Nearest Neighbor (WNN) analysis separated C10 and SVEC cells, exemplified by the concordant HMGB2 protein and *Hmgb2* RNA expression patterns (Fig. 2k). Multi-Omics Factor Analysis (MOFA) identified latent factors that explained different proportions of variance across the proteomic and transcriptomic modalities (Fig. 2l); for instance, Factor 1 explained variance in both modalities but was associated with largely distinct high-weight protein and RNA features (Fig. 2m). Taken together, these results show that msmu-processed proteomics data can be reliably reused and readily integrated with other modalities.

In summary, the modular design of msmu and architectural advantages gained from using MuData facilitate flexible proteomics data processing and multimodal interoperability. Importantly, msmu embeds machine-actionable provenance directly within the proteomics data object, allowing recorded workflows to be inspected, verified, and replayed. Exported workflow scripts can be modified to create alternative branches from the original analysis, supporting transparent parameter exploration and reuse. Future development will broaden search engine support and the range of preprocessing and statistical methods. By making machine-actionable provenance a first-class component of the data object, msmu provides a foundation for FAIR-aligned analysis of proteomics data.

## Methods

### Technical specification of msmu

msmu requires Python ≥ 3.12. Core dependencies include MuData, AnnData, polars, sdrf-pipelines, statsmodels, scipy, scikit-learn, umap-learn, directlfq, inmoose, pyopenms, plotly, and fastparquet; complete dependencies and version requirements are listed in pyproject.toml.

### Provenance recording and workflow replay

Supported msmu operations that transform MuData are instrumented to record provenance in MuData.uns[“_log”]. For each successfully recorded call, msmu stores the function name and qualified path, parameters including defaults and input descriptors, execution times, parent event identifiers, output descriptors, and an execution-environment identifier. When hashing is enabled, input and output states are hashed using SHA-256 under a fixed floating-point normalization policy, which normalizes floating-point values to 12 significant digits for hashing while leaving the stored values and numerical calculations unchanged. Before executing a provenance-supported function, msmu compares the input hash with the output hash from the previously recorded step. A mismatch triggers a warning that the data have changed outside the recorded workflow. Execution-environment records include the Python version, operating system, architecture, and installed package versions.

mm.pv.to_env(), mm.pv.to_script(), and mm.pv.replay() take a MuData object, or its log decoded with mm.pv.get_log(), as input and leave it unchanged. mm.pv.to_env() exports the recorded Python version and exact package pins in requirements.txt format for uv or environment.yml format for conda, with the recorded platform and msmu source revision as comments; if events reference differing environments, one must be selected. These are version pins rather than a lockfile, so package sources, builds, and native libraries are not reconstructed.

mm.pv.replay() and mm.pv.to_script() share a planning step that orders recorded events topologically, resolves function paths only among public msmu functions and checks recorded parameters against current function signatures. Replayable histories start from recorded file readers and consist of operations that return a MuData object, including branches and merges; histories containing operations with separate result objects, such as DE analysis, cannot be replayed. Histories with unrecorded modifications, detected as mismatches between an event’s recorded input hash and its parent’s output hash, are refused. mm.pv.replay() first compares the current execution environment with the recorded one and reports differences as warnings without installing packages. It then reruns the events from the source input files at their recorded paths, under each event’s recorded hash normalization policy, checking the hashes of recorded input files and incoming data states before each step and of the regenerated output after it, and stops at the first mismatch; verify=False skips these checks. Replay returns a new MuData object whose provenance log records the replayed events and the current environment. mm.pv.to_script() writes the same plan as a Python script without reading inputs or executing operations, rendering each event as a public msmu call with all recorded parameters, including defaults, as keyword arguments. By default (verify=True), the script embeds the recorded hashes and performs the same checks as replay. With verify=False, only the calls are retained, so parameters, input files, or steps can be edited for customized reuse without halting on hash mismatches. In both modes, the script checks the environment and enables hashing, so every msmu operation it runs is recorded in a new provenance log; changes made outside msmu functions are flagged by the continuity check.

### Database searching

Peptide identification of the Fulcher *et al*. dataset^23^ was performed using Sage. *In silico* digestion was set to fully tryptic, allowing cleavage after lysine/arginine but restricting before proline. Fragment ion *m/z* ranges were set to 150–2,000, and peptide mass ranges were set to 500–5,000 Da. Carbamidomethylation of cysteine was specified as a fixed modification, while methionine oxidation and protein N-terminal acetylation were set as variable modifications. Label-free quantification was performed using FlashLFQ, with match-between-runs disabled and quantification restricted to PSMs with a false discovery rate < 1%.

### Processing of the single-cell proteomics data

The re-searched proteomics data were processed using msmu and stored in MuData format. First, identification and quantification data generated by Sage and FlashLFQ were ingested. PSMs passing q < 0.01 were aggregated to the peptide level and matched to FlashLFQ-derived peptide intensities. Peptides that did not pass the q-value cutoff (< 0.01) were removed, followed by log_2_ transformation and pairwise-median normalization of peptide intensities. Protein-level data were obtained by summarizing peptide-level data using a parsimony-based protein inference procedure and Tukey’s median polish method for intensity aggregation. A protein-level q-value cutoff (< 0.01) was also applied. To support downstream analysis requiring complete observations, the PIMMS^26^ package was used to impute missing values. Missing-value pattern assessment and robust missing-value imputation were applied sequentially, and the resulting matrix was stored as an additional layer at the protein-level AnnData. The processed MuData object was written as an .h5mu file.

### Processing of the single-cell transcriptomics data

Raw count matrices were processed using scanpy. All files were imported and merged into one AnnData object, followed by filtering of low-quality cells and genes with low expressions. The matrix was then normalized at the cell level, and expression values were log-transformed. The processed AnnData object was written as an .h5ad file.

### Integrated analysis of the single-cell dataset

The proteomics .h5mu and transcriptomics .h5ad files were loaded as MuData and AnnData objects, respectively. The transcriptomics AnnData was added to the proteomics MuData object as an RNA modality. Prior to integration, cells (.obs) were aligned to share the same identifiers across modalities. WNN Uniform Manifold Approximation and Projection (UMAP) coordinates were calculated with scanpy and muon from the scverse ecosystem. PCA and neighborhood graphs were computed for both the protein and RNA modalities. Multimodal neighbors were then identified using a WNN scheme. The resulting graph was embedded using UMAP to generate WNN UMAP coordinates. MOFA was additionally performed using the muon implementation, which created an HDF5 model file for downstream factor analysis.

### Cross-platform reproducibility assessment

The reference workflow was executed on macOS 27.0.1 (arm64) and Debian 13.6 (x86_64) using msmu version 0.4.3. The exported environment specification was used to create a Python 3.13.5 environment with matched Python and package versions, and the workflow was replayed from the same source input files with hash verification enabled. Reproducibility of protein inference was assessed with Jaccard indices for peptide nodes, protein nodes, and peptide–protein edges of the protein inference map. Numerical reproducibility was quantified as the RMSD and max |Δ| between paired _−_log_10_ protein-level q-values (n = 933 proteins), log_2_ protein intensities (n = 23,238 non-missing protein–cell values), and _−_log_10_ DE q-values. For DE, reproducibility was also assessed by comparing significance calls at q < 0.10, 0.05, and 0.01.

## Data availability

The MS data from Fulcher *et al*.^23^ reanalyzed in this study are available from the MassIVE repository under accession MSV000089280. The corresponding scRNA-seq data can be downloaded from the Gene Expression Omnibus (GEO)^27^ under accession GSE201575. The .h5mu file created using msmu for the single-cell multi-omics dataset and a companion tutorial detailing how to extract quantitative matrices, annotations, and provenance associated with the processing workflow can be found at https://bertis-informatics.github.io/msmu/tutorials/2024_Fulcher/.

## Code availability

A Python package for msmu is released via PyPI, and its source code is available at the GitHub repository: https://github.com/bertis-informatics/msmu. All scripts used for the analyses reported in this study are also provided in the repository. The full documentation, including tutorials and API references, can be found at https://bertis-informatics.github.io/msmu/. msmu is distributed under the BSD 3-Clause License.

## Acknowledgements

This work was supported by a National Research Foundation of Korea (NRF) grant funded by the Korea government (MSIT) (RS-2024-00454407). We thank Jungkap Park for providing feedback on the manuscript.

## Author information

These authors contributed equally: Hyung-Wook Choi, Byeongchan Lee.

### Contributions

H.-W.C. and B.L. conceived the method and developed the msmu package. S.H. gave feedback on the

method and results. H.-W.C., B.L., and S.H. wrote the paper. U.-B.K. gave feedback for the paper. All authors read and approved the paper.

