## Supplementary Table 1 for "msmu: a computational framework for provenance-aware mass spectrometry-based proteomics analysis"

### Supplementary Information

#### Supplementary Table 1. Comparison of msmu with AnnData- and MuData-based software for post-search proteomics data analysis.

| | Capability | msmu (this work) v0.4.3 | AlphaPeptTools[1] v0.4.0 | ProteoPy[2] v0.1.1 | scpviz[3] v1.1.2 | grassp[4] v0.6.0 |
|---|---|---|---|---|---|---|
| **Data Model and Interoperability** | | | | | | |
| 1 | One object holding PSM/precursor, peptide and protein levels | ● MuData with one AnnData modality per level (PSM/ precursor, peptide, protein, PTM site); .h5mu | ● AnnData per level (precursor, peptide, protein, gene) in MuData; .h5ad, .h5mu | ○ Separate peptide- or protein-level AnnData objects; .h5ad | ◐ pAnnData class wrapping peptide and protein AnnData; no native file format (CSV export) | ○ Protein-level AnnData with proteins as observations; .h5ad |
| 2 | Queryable links between features across levels | ● Peptide–protein inference map in .uns | ● Feature-mapping adjacency in .varp with ancestor and descendant queries | ◐ Protein identifiers in peptide-level .var; no links between objects | ● Protein–peptide relationship matrix (.rs) | ○ |
| 3 | Native AnnData/MuData model for scverse multimodal analysis | ● AnnData and MuData | ● AnnData and MuData | ● AnnData | ◐ AnnData per level inside the custom class; scanpy, Harmony | ● AnnData; built on scanpy |
| **Input and Sample Metadata** | | | | | | |
| 4 | Supported search engines | ● Sage, DIA-NN, MaxQuant, FragPipe, Delpi (PSM/precursor tables) | ● Sage, AlphaDIA, DIA-NN, MaxQuant, FragPipe, Spectronaut, pFind (PSM/ precursor tables); AlphaPept, Proteome Discoverer, mzTab (protein-group tables) | ◐ DIA-NN; generic long-format tables | ◐ DIA-NN, Proteome Discoverer (peptide and protein tables) | ○ Protein-group matrices from MaxQuant, FragPipe, DIA-NN |
| 5 | SDRF-Proteomics sample metadata | ● Validated with sdrf-pipelines | ○ | ○ | ○ | ○ |
| 6 | Label-free DDA, DIA, and isobaric (TMT) quantification | ● LFQ, DIA, TMT | ◐ LFQ, DIA, TMT | ◐ LFQ, DIA | ◐ LFQ, DIA | ◐ LFQ, DIA, spectral counts |
| **Post-Search Preprocessing** | | | | | | |
| 7 | Peptide- and protein-level target–decoy q-value estimation | ● Target–decoy q-values at peptide and protein levels | ○ Taken from the search engine | ○ | ○ Taken from the search engine | ○ |
| 8 | Protein inference | ● Parsimony-based protein inference | ○ Upstream protein groups | ○ None (COPF proteoform analysis) | ○ Upstream protein groups | ○ Upstream protein groups |
| 9 | Summarization from PSM/precursor to protein level | ● PSM→peptide: median, mean, sum, top-N. Peptide→protein: median, mean, sum, top-N, Tukey median polish, directLFQ | ○ None; per-level values from the search engine | ● Peptide→protein: sum, mean, median, maximum, custom function | ◐ Peptide→protein by directLFQ only | ○ None; protein-group input |
| 10 | Normalization | ● Median, median centering, quantile, total sum, pairwise median | ● Total mean or median, IRS, scale-and-center | ◐ Median | ● Sum, median, mean, maximum, reference feature, robust scaling, quantile transform, directLFQ | ◐ Total-intensity normalization |
| 11 | Batch correction | ● IRS, median centering, ComBat; LOWESS for continuous covariates | ◐ ComBat | ◐ Median normalization within batches | ◐ Harmony on the PCA embedding only | ○ |
| 12 | Missing-value imputation | ○ None; user-imputed matrices can be stored as additional layers. | ● Down-shifted Gaussian, kNN, median, BPCA | ● Down-shifted Gaussian | ● Mean, median, minimum, kNN, PIMMS (DAE, VAE, CF) | ● Down-shifted Gaussian, kNN |
| 13 | PTM-site quantities adjusted for parent-protein abundance | ● Ratio or ridge adjustment | ○ | ○ | ○ | ○ |
| **Statistics and Visualization** | | | | | | |
| 14 | Differential expression with linear models and covariates | ● limma moderated t with covariates; permutation tests (Welch, Student, Wilcoxon); BH or empirical FDR | ● Welch t; moderated t with a categorical covariate; AlphaQuant; BH | ◐ t, Welch t, one-way ANOVA; BH or Bonferroni | ● t, Welch t, Mann–Whitney, Wilcoxon; mixed models or pseudobulk with covariates; BH | ◐ t-test of enrichment against untagged controls or other fractions |

| | Capability | msmu (this work) v0.4.3 | AlphaPeptTools[1] v0.4.0 | ProteoPy[2] v0.1.1 | scpviz[3] v1.1.2 | grassp[4] v0.6.0 |
|---|---|---|---|---|---|---|
| 15 | Dimensionality reduction (PCA and UMAP) | ● PCA, UMAP | ◐ PCA | ◐ PCA, UMAP through scanpy (tutorials) | ● PCA, UMAP, Leiden clustering | ● PCA, UMAP, aligned UMAP across datasets |
| 16 | Built-in QC metrics and plots | ● Identification counts, intensity distributions, completeness, correlation, UpSet, precursor purity | ● CV, MAD, completeness, total intensity, PC regression | ● Completeness, CV, correlation, intensity distributions | ● CV, abundance rank, raincloud, correlation, clustermap; STRING and GSEA enrichment | ● QC metrics, replicate CV, highly variable proteins, marker profiles |
| **Provenance and Reproducibility** | | | | | | |
| 17 | Processing history stored in the data object | ● Provenance log in MuData .uns, saved in the .h5mu file | ○ | ○ | ◐ Time-stamped free-text history held in memory | ○ |
| 18 | Step parameters, software versions, execution environment recorded in machine-readable form | ● Parameters, including defaults, for every recorded call, Python, OS, architecture, package versions | ○ | ◐ Result keys encode the test, grouping and design | ○ | ○ |
| 19 | Content hashes of input and output data states | ● SHA-256 at 12 significant digits | ○ | ○ | ○ | ○ |
| 20 | Detection of unrecorded modifications before each step | ● Hash continuity check | ○ | ○ | ○ Flags its derived summary table as stale when that table is edited outside the workflow; the data matrices are not checked. | ○ |
| 21 | Export of an executable analysis script | ● mm.pv.to_script() | ○ | ○ | ○ | ○ |
| 22 | Export of an environment specification | ● mm.pv.to_env() | ○ | ○ | ○ | ○ |
| 23 | Replay of the recorded workflow from the saved object | ● mm.pv.replay() | ○ | ○ | ○ | ○ |

● built in and documented ◐ partial: limited, indirect, or recorded outside the data object ○ absent, or not found in the documentation or source code
